# Generation and Characterization of a Knockout Mouse of an Enhancer of *EBF3*

**DOI:** 10.1101/2025.01.09.631762

**Authors:** Emily Cordova Hurtado, Janine M. Wotton, Alexander Gulka, Crystal Burke, Jeffrey K. Ng, Ibrahim Bah, Juana Manuel, Hillary Heins, Stephen A. Murray, David U. Gorkin, Jacqueline K. White, Kevin A. Peterson, Tychele N. Turner

**Affiliations:** Department of Genetics, Washington University School of Medicine, St. Louis, MO 63110, USA; The Jackson Laboratory, Bar Harbor, ME, 04609, USA; Department of Biology, Emory University. Atlanta, GA 30322, USA

**Author notes:** co-first authors (Emily Cordova Hurtado, Janine M. Wotton). Co-Correspondence to,; Tychele N. Turner, Ph.D., Assistant Professor, Washington University School of Medicine, Department of Genetics, 4523 Clayton Avenue, Campus Box 8232, St. Louis, MO 63110; Kevin Peterson, Ph.D., Senior Research Scientist, Technology & Resource Development, The Jackson Laboratory, Bar Harbor, ME, USA; Jacqueline White Ph.D., Senior Director - Center for Biometric Analysis, Deputy Director - Scientific Services JAX MG, The Jackson Laboratory, 600 Main Street, Bar Harbor, ME 04609.

## Abstract

Genomic studies of autism and other neurodevelopmental disorders have identified several relevant protein-coding and noncoding variants. One gene with an excess of protein-coding *de novo* variants is *EBF3* that also is the gene underlying the Hypotonia, Ataxia, and Delayed Development Syndrome (HADDS). In previous work, we have identified noncoding *de novo* variants in an enhancer of *EBF3* called hs737 and further showed that there was an enrichment of deletions of this enhancer in individuals with neurodevelopmental disorders. In this present study, we generated a novel mouse line that deletes the highly conserved, orthologous mouse region of hs737 within the Rr169617 regulatory region, and characterized the molecular and phenotypic aspects of this mouse model. This line contains a 1,160 bp deletion within Rr169617 and through heterozygous crosses we found a deviation from Mendelian expectation (p = 0.02) with a significant depletion of the deletion allele (p = 5.8 × 10^-4^). *Rr169617^+/-^* mice had a reduction of *Ebf3* expression by 10% and *Rr169617^-/-^* mice had a reduction of *Ebf3* expression by 20%. Differential expression analyses in E12.5 forebrain, midbrain, and hindbrain in *Rr169617^+/+^*versus *Rr169617^-/-^* mice identified dysregulated genes including histone genes (i.e., *Hist1h1e*, *Hist1h2bk*, *Hist1h3i*, *Hist1h2ao)* and other brain development related genes (e.g., *Chd5*, *Ntng1*). *A priori* phenotyping analysis (open field, hole board and light/dark transition) identified sex-specific differences in behavioral traits when comparing *Rr169617^-/-^* males versus females; whereby, males were observed to be less mobile, move slower, and spend more time in the dark. Furthermore, both sexes when homozygous for the enhancer deletion displayed body composition differences when compared to wild-type mice. Overall, we show that deletion within Rr169617 reduces the expression of *Ebf3* and results in phenotypic outcomes consistent with potential sex specific behavioral differences. This enhancer deletion line provides a valuable resource for others interested in noncoding regions in neurodevelopmental disorders and/or those interested in the gene regulatory network downstream of *Ebf3*.

## INTRODUCTION

Autism is a neurodevelopmental disorder with high heritability ^1; 2^. Several studies focusing on exome sequencing have identified *de novo* variants (DNVs) that disrupt genes ^3-13^. Other genetic factors include large copy number variants ^8; 14-22^ and common variants contributing to polygenic risk ^23^, respectively. A contribution from noncoding DNVs has also been identified from studies using whole-genome sequencing ^24-32^. We previously identified an enhancer, hs737, with an excess of noncoding DNVs in individuals with autism ^33^. This enhancer targets the gene *EBF3* that is the underlying gene for Hypotonia, Ataxia, and Delayed Development Syndrome (HADDS). Protein-coding DNVs of *EBF3* are also known to be genome-wide significant for excess in neurodevelopmental disorders ^33-37^. When comparing individuals with protein-coding DNVs in *EBF3* to those with noncoding DNVs in hs737, that affects *EBF3*, we found that individuals with protein-coding DNVs are more severe in their phenotype ^33^. Beyond single point variants in this enhancer, we also previously showed that it does not deviate from the copy number of two in 56,256 alleles from individuals who do not have neurodevelopmental disorders^33^. However, it is enriched for deletions and nominally enriched for duplications in individuals with neurodevelopmental disorders ^33^.

The *EBF3* gene encodes a transcription factor that preferentially binds to the promoters of other transcription factors and chromatin-binding proteins involved in neurodevelopmental disorders (NDDs) (e.g., *CHD2*, *CHD8*, *ARID1B*) ^33^. This gene is a member of the EBF gene family, which includes EBF1, EBF2, EBF3, and EBF4 ^38^, and is known to form homodimers or heterodimers with itself or other family members, respectively. It is known to be regulated by the X chromosome gene *ARX* that is also involved in NDDs. It resides in a large TAD region in the genome of ∼2 Mbp and several regulatory regions of *EBF3* exist within the TAD. The hs737 enhancer is ∼1.5 Mbp from the promoter of *EBF3* and has been shown to contact the promoter ^33; 39^. It is an enhancer that is a member of the VISTA enhancer database that contains several enhancers with conservation in human, mouse, and rat ^40^. While expression of *EBF3* is ubiquitous in the human body, the activity of hs737 seems to be restricted to the fetal brain ^33^.

As noted, there is an enrichment of DNVs within hs737 in individuals with autism and an enrichment of deletions in individuals with neurodevelopmental disorders. We sought to determine the molecular and phenotypic consequence of deletion of hs737 in a model system. Thus, we focused on generating a mouse model for this genomic interval as the sequence of hs737 is highly conserved with its orthologous mouse sequence (within the Rr169617 https://www.informatics.jax.org/marker/MGI:7057839 regulatory region) ^33^. Here, we describe the creation of a novel mouse line engineered to delete the relevant sequence within Rr169617, assess the molecular consequences through RNAseq experiments, and determine the phenotypic consequences through systematic broad-based phenotyping assays. This mouse model provides a useful tool to others in the field especially those studying the *EBF3* gene regulatory network (GRN) that has been implicated in autism and other neurodevelopmental disorders ^33-37^.

## MATERIALS AND METHODS

### Generation of Deletion Mouse Lines

To delete the relevant sequence in Rr169617, paired upstream (CATGCAGAGAAAACAAAATG, GCTGAATTGTAGCGTGTTTA) and downstream (TGGCGCCAGTGGGCCCCGAC, ATCCTGGCACTGGCGCCAGT) guides were identified to flank the genomic region of interest on mouse chr7:136083335-136084349 (GRCm38/mm10). Guide RNAs were incubated with Cas9 protein to generate ribonucleoprotein complexes (RNPs) followed by electroporation into C57BL/6J zygotes (JAX strain #:000664) using standard conditions. Following PCR genotyping for the deletion allele three independent founder lines (lines 299, 300 and 304) were recovered and backcrossed to C57BL/6J to generate N1 progeny; however, only two lines (299 and 304) showed successful germline transmission. A molecular description of the genomic lesion present in each of these independent lines was defined by Sanger Sequencing of PCR amplicons. Line 299 carried a 1,160 bp deletion (chr7:136083275-136084434 GRCm38/mm10); referred to as C57BL/6J-Rr169617^em1Tnt^/J (MMRRC # to be added upon acceptance of the line to the database). Line 304 was found to contain a 1,147 bp deletion (chr7:136083283-136084428 GRCm38/mm10); referred to as C57BL/6J-Rr169617^em2Tnt^/J (MMRRC # to be added upon acceptance of the line to the database). Both lines have been cryopreserved and will be publicly available from The Jackson Laboratory Mutant Mouse Resource and Research Center. In this study, detailed characterization was performed on C57BL/6J-Rr169617^em1Tnt^/J that we will refer to as Rr169617 throughout the rest of this study.

### Ethical Approval

All mouse work reported herein was conducted at the Jackson Laboratory under the Institutional Animal Care and Use Committee-approved license numbers 11005 and 20028. AAALACi accreditation number 00096, and NIH Office of Laboratory Animal Welfare assurance number D16-00170.

### Animal Housing Information

Animals used in phenotyping studies were homozygous mutant *Rr169617^-/-^*mice [female (n=8), male (n=8)] and age and sex matched wildtype control *Rr169617^+/+^*mice [female (n=9), male (n=10)]. Mice were housed (1 to 5 animals per cage) in individually ventilated cages [Thoren Duplex II Mouse Cage #11 and Thoren Maxi-Miser PIV System (30.8 L x 30.8 W x 16.2 H cm)] behind a pathogen-free barrier. Access to water and food (5K52 diet, LabDiet) was *ad libitum*. Wood shavings (aspen) bedding substrate was provided and sections housing individual mice were supplemented with environmental enrichments (e.g. a nestlet and cardboard hut). Mice were housed in rooms with 12-hour light–dark cycle and temperature and humidity were maintained between 20-22°C and 44-60%, respectively.

### Mouse Colony Maintenance and Embryo Collections

Mouse colonies were maintained by either backcrossing to wild-type C57BL/6J or by intercrosses between heterozygous animals. Timed matings were performed by intercrossing heterozygous *Rr169617^+/-^* animals where noon of the day of detection of vaginal plug was considered embryonic day 0.5 (E0.5). Embryos were kept cold on ice in 1X phosphate buffered saline (PBS) and microdissected in ice cold PBS. Embryonic tissues were snap frozen in liquid nitrogen and stored at -80°C until use.

### Genotyping PCR for the Deletion

DNA was extracted from E12.5 forebrains for one sample each of wild type (*Rr169617^+/+^*), heterozygous (*Rr169617^+/-^*), and homozygous (*Rr169617^-/-^*) mice using the Zymo Quick-DNA HMW MagBead kit. This extraction method was also used to derive DNA from the HT-22 cell line as a control for the PCR. Primers were designed to test for presence of the deletion in Rr169617 (mm10 chr7:136083275-136084434). The forward primer was 5’ CATACTTAGCTACTGTGGATGGTGA 3’ and the reverse primer was 5’ CAAATCCCACCTTAACAGCACATAG 3’. PCR reactions consisted of 30 ng HMW DNA of each sample, positive control (HT-22), or negative control (water), 1.25 µl of 10 µM forward primer, 1.25 µl of 10 µM reverse primer, 0.75 µl DMSO, 12.5 µl 2× Phusion High Fidelity master mix, and nuclease free water up to 25 µl. Cycling conditions were 98°C for 2 minutes, 25 cycles of [98°C for 10 seconds, 70°C for 30 seconds, 72°C for 30 seconds], 72°C for 10 minutes, 4°C hold. The samples were run on an Agilent Bioanalyzer. The wild-type band was 2,618 bp and the deletion-containing band was 1,484 bp. Confirmation of the sequence of the wild type and deletion bands were completed by TOPO TA cloning of the sequences into a plasmid (using the TOPO TA Cloning Kit for Sequencing) and sequencing of the plasmid by Oxford Nanopore Technology sequencing at Plasmidsaurus.

### Long-Read Whole-Genome Sequencing of Mice

E12.5 mouse forebrains were pooled from three wild type (*Rr169617^+/+^*) and three homozygous mice (*Rr169617^-/-^*), respectively. High molecular weight DNA was extracted for each pooled sample. Each pool was made into a library for PacBio HiFi sequencing on the Revio sequencer. Each library was sequenced using one SMRT cell to approximately 30× coverage.

### RNA extraction, cDNA synthesis and RNA-seq of E12.5 Forebrain

Mouse E12.5 fetal forebrain tissue from mice that were wild-type (*Rr169617^+/+^*) (n = 12), heterozygous (*Rr169617^+/-^*) (n = 14), or homozygous (*Rr169617^-/-^*) (n = 10) for the deletion in Rr169617 was used to extract RNA. The RNA was extracted using a Bead Bug homogenizer to homogenate the tissue and the Maxwell simplyRNA Tissue kit for RNA extraction. SuperScript III First-Strand Synthesis System was used for reverse transcription. Taqman mouse *Ebf3* (Mm00438642_m1) and GAPDH (Mm99999915_g1) gene expression assays were performed on a QuantStudio 6 Flex quantitative thermocycler using four reactions for each sample. QuantStudio Real-Time PCR software was used to run the thermocycler using the Standard Comparative Ct (ΔΔCt) method. Three individuals performed a total of five qPCR assays in quadruplicate. Results were reviewed by three individuals to assess each set of quadruplicates for outliers (>0.5 cycles apart), and these were removed from the data sets. For RNA-seq, three RNA samples from each E12.5 genotype group (*Rr169617^+/+^*, *Rr169617^+/-^*, *Rr169617^-/-^*) were polyA selected and sequenced to a target of 200 million read pairs using Illumina NovaSeq6000. Every sample RNA had a RIN greater than 8.0. Ribosomal RNA was removed through poly-A selection with Oligo-dT beads (mRNA Direct kit, Life Technologies). The mRNA was fragmented in reverse transcriptase buffer and heated to 94°C for 8 minutes. Reverse transcription of the mRNA to cDNA was performed using the SuperScript III RT enzyme with random hexamers. A second strand synthesis was carried out to produce double-stranded cDNA. The cDNA was blunt-ended, an A base was added to the 3’ ends, and Illumina sequencing adapters were ligated to the ends. The ligated fragments were amplified for 12-15 cycles with primers incorporating unique dual index tags. Finally, the fragments were sequenced on an Illumina NovaSeq with paired-end reads extending 150 bases at the McDonnell Genome Institute.

### RNA-seq of E12.5 Midbrain and Hindbrain

RNA was extracted from E12.5 midbrains and hindbrains of three independent samples of each genotype (*Rr169617^+/+^*, *Rr169617^-/-^*), respectively. Library preparation, ribosomal RNA reduction, and Illumina UDI library preparation were performed at the University of Maryland Institute for Genome Sciences. They were sequenced (rRNA depletion RNAseq) to a target of 200 million read pairs using an Illumina NovaSeq6000.

### RNA-Seq Analysis

The RNA-seq analysis was run using the ENCODE pipeline, found here (https://github.com/ENCODE-DCC/rna-seq-pipeline), due to it being a well-developed standard. The only modifications were hardcoded PATH variables so that the pipeline would function properly on our HPC. The mouse Gencode M21 reference data was used; links are provided in the ENCODE documentation. The forebrain poly-A samples were run as paired, unstranded runs, while the rRNA-depleted samples were run as paired, reverse-stranded runs. Differential gene expression analysis was performed using DESeq2 ^41^.

### Phenotype Pipeline

The mice progressed through the JAX KOMP phenotyping pipeline (Supplementary Table 5, https://www.mousephenotype.org/impress/PipelineInfo?id=12)^42^. The methods for all assays in the pipeline are provided online (https://www.mousephenotype.org/impress/PipelineInfo?id=12) and the assays for which we had *a priori* hypotheses are detailed below.

### Behavioral Assays

Three behavioral assays (open field, light/dark transition and hole board) were conducted to provide information on anxiety, exploration and mobility. Testing was conducted between 7am and 5pm in the light portion of their 24-hour cycle and mice were first habituated to the room for 30 minutes. For all three assays, mice were placed in an acrylic chamber (40 x 40 x 40 cm) contained within a sound attenuated, ventilated cabinet (64 L x 60 W x 60 H cm) and the motor behavior and location were recorded by horizontal infrared photobeam sensors (16 x 16 array) using Fusion behavioral tracking software (Omnitech Electronics, Columbus, OH, USA).

At approximately 8 weeks of age mice completed the open field test (OF). Mice were placed in the center of the open field arena (light level:100-200 lux) and behavior was recorded for 20 minutes. The following week mice were tested in the light/dark transition assay (L/D) which included a dark insert chamber (40 x 20 x 40 cm) so that half the chamber was in light (∼200 lux) and half dark (∼1 lux). Mice were placed in the lit portion of the chamber facing the dark portion and were recorded for 20 minutes. Later the same week mice were tested in the hole board (HB) configuration which included a grid of 16 shallow holes in the floor (4 x 4 grid) of the open field arena for the mice to explore (light level:100-200 lux) and behavior was recorded for 10 minutes.

### Dual-Energy X-ray Absorptiometry (DEXA)

DEXA provided measures of body composition (lean and fat mass), bone mineral content and bone density. Mice (approximately 14 weeks of age) were anesthetized [intraperitoneal injection of 400 mg/kg tribromoethanol diluted in sterile PBS (in-house pharmacy)], measured for length and then placed in the previously calibrated densitometry machine (Lunar Piximus II from GE Medical systems). The region of interest measured excluded the head and neck.

### Statistical Analysis of Phenotypes

Targeted *a priori* hypotheses: Although the mice were tested in a full phenotyping pipeline, several of the assays provided measures that were hypothesized to differ between the sexes of the mutant line ^33^. These *a priori* hypotheses were analyzed first. The three described behavior tests (OF, L/D, HB) were conceptual replications using the same equipment in different configurations. The data was therefore analyzed using repeated measures ANOVA for the common behavioral parameters using the between-subject’s factors of sex (male, female), genotype (*Rr169617^-/-^*, *Rr169617^+/+^*) and interaction between sex and genotype. Males were predicted to be more affected ^33^ therefore planned comparisons of strain were completed for each sex for each assay with Bonferroni adjustment for multiple testing for each analysis. Tests of anxiety were treated as independent multivariate ANOVAs for each relevant parameter.

The body composition (DEXA) data was analyzed as a multivariate ANOVA with factors of sex (male, female), genotype (*Rr169617^-/-^*, *Rr169617^+/+^*) and interaction between sex and genotype. Planned comparisons of strain were completed for each sex for each assay with Bonferroni adjustment for multiple testing. The *a priori* hypotheses were analyzed using SPSS version 29 (IBM).

All parameters: The phenotyping pipeline was analyzed using standard IMPC analysis based on PhensStat ^43^ which was designed to find the best analysis for high throughput data. Standard mandatory parameters for the IMPC were analyzed (Supplementary Table 5, https://www.mousephenotype.org/impress/PipelineInfo?id=12). Continuous data were analyzed using optimized linear model ANOVAs with the initial factors of genotype, sex and bodyweight when available. Categorical data (e.g. eye morphology) were tested using Fisher’s Exact tests.

## RESULTS

### The Topologically Associating Domain with Hs737 is Highly Conserved in Mouse

By aggregating genomic annotation data from ours ^33^ and others previous work ^39; 44; 45^, we studied the features of the human genomic topologically associating domain (TAD) region containing hs737 in the mouse genome (within Rr169617). EBF3 resides within TAD1949 originally defined in Dixon et al. 2012 ^44^ via Hi-C in human embryonic stem cells. To identify the orthologous region in mouse, we performed *liftover* from human (GRCh38/hg38) to mouse (GRCm38/mm10) and found that this TAD was mostly conserved in mouse (**Figure 1**). The large TAD region contains the same TAD boundaries as seen in human (**Figure 1**). Further support for the conserved architecture of this region is provided by comprehensive capture-Hi-C experiments for several VISTA enhancers that identified multiple enhancer-promoter interactions between the region within Rr169617 and the promoter of *Ebf3* (**Figure 1**) ^39^ These findings are consistent with our previous observations of hs737 contacting the promoter of *EBF3* by examining Hi-C data in the human fetal brain ^33; 39^.

**Figure 1:**
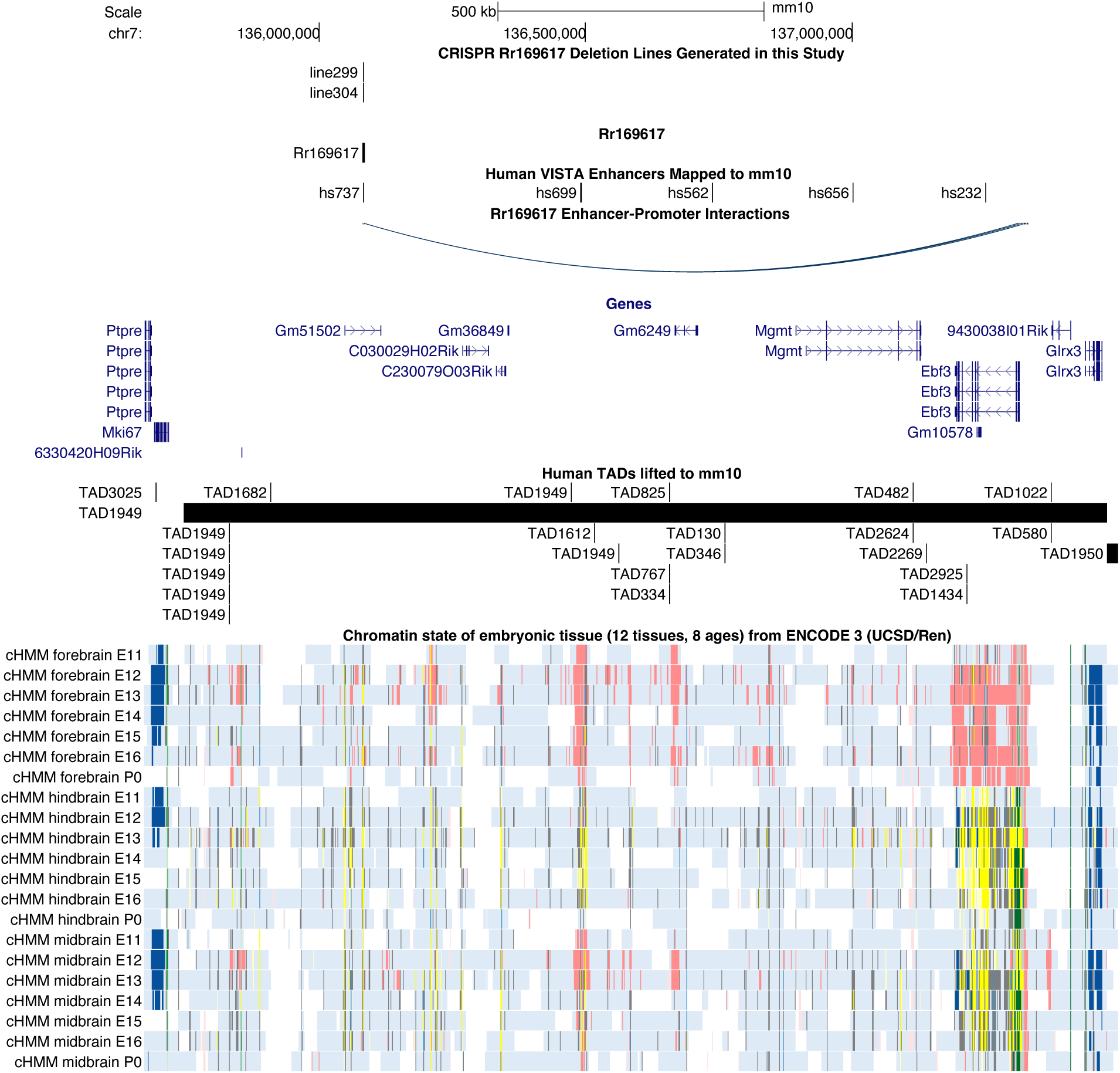
*Ebf3* regulatory landscape and associated hs737/Rr169617 enhancer deletion mouse lines. Genome browser view of the topologically associating domain region containing Rr169617 and its target gene *Ebf3* (GRCm38/mm10). The first track shows the two independent founder mouse lines generated in this study: line 299 (C57BL/6J-Rr169617^em1Tnt^/J) and line 304 (C57BL/6J-Rr169617^em2Tnt^/J). The second track shows the location of the regulatory region, Rr169617. The third track shows the location of human VISTA enhancers lifted over to the mouse genome. Included is hs737 that resides within the Rr169617 region. The fourth track shows enhancer-promoter interactions of Rr169617 and *Ebf3* from Chen et al. 2024, *Nature Genetics*. The fifth track shows the genes within the region. The fifth track shows human topologically associating domains lifted over to this region and show high conservation. Finally, the chromatin state data available from ENCODE3 is shown across the different timepoints in mouse development.

### Generation of Knockout Mouse Lines

A deletion within Rr169617 was generated in mouse using CRISPR/Cas9 and paired guide RNAs flanking the region of interest. This resulted in three separate deletion founder animals (**Figure 1**). Due to the similarity in the deletions, detailed experimental characterization was performed on line 299 that has a 1,160 bp deletion (chr7:136083275-136084434 GRCm38/mm10). To further validate this line, we performed PacBio HiFi long-read whole-genome sequencing on E12.5 forebrain tissue from wild type (*Rr169617^+/+^*) and homozygous deletion animals (*Rr169617^-/-^*). All of the sequence reads in the homozygous deletion (*Rr169617^-/-^*) mice contained a 1,160 bp deletion and none of the sequence reads in the wild type (*Rr169617^+/+^*) mice contained the deletion (**Figure 2A**). Since we performed long-read sequencing, we also assessed the methylation (5mC) status within the enhancer region (**Figure 2B**) and over the *Ebf3* promoter (**Supplementary Figure S1**). The sequence deleted in *Rr169617^-/-^* contained CpG sites that were not methylated in *Rr169617^+/+^*. The methylation status surrounding the deletion region was similar in both *Rr169617^+/+^*and *Rr169617^-/-^* (**Figure 2B**). The *Ebf3* promoter sequence was mostly not methylated in both *Rr169617^+/+^* and *Rr169617^-/-^* (**Supplementary Figure S1**).

**Figure 2:**
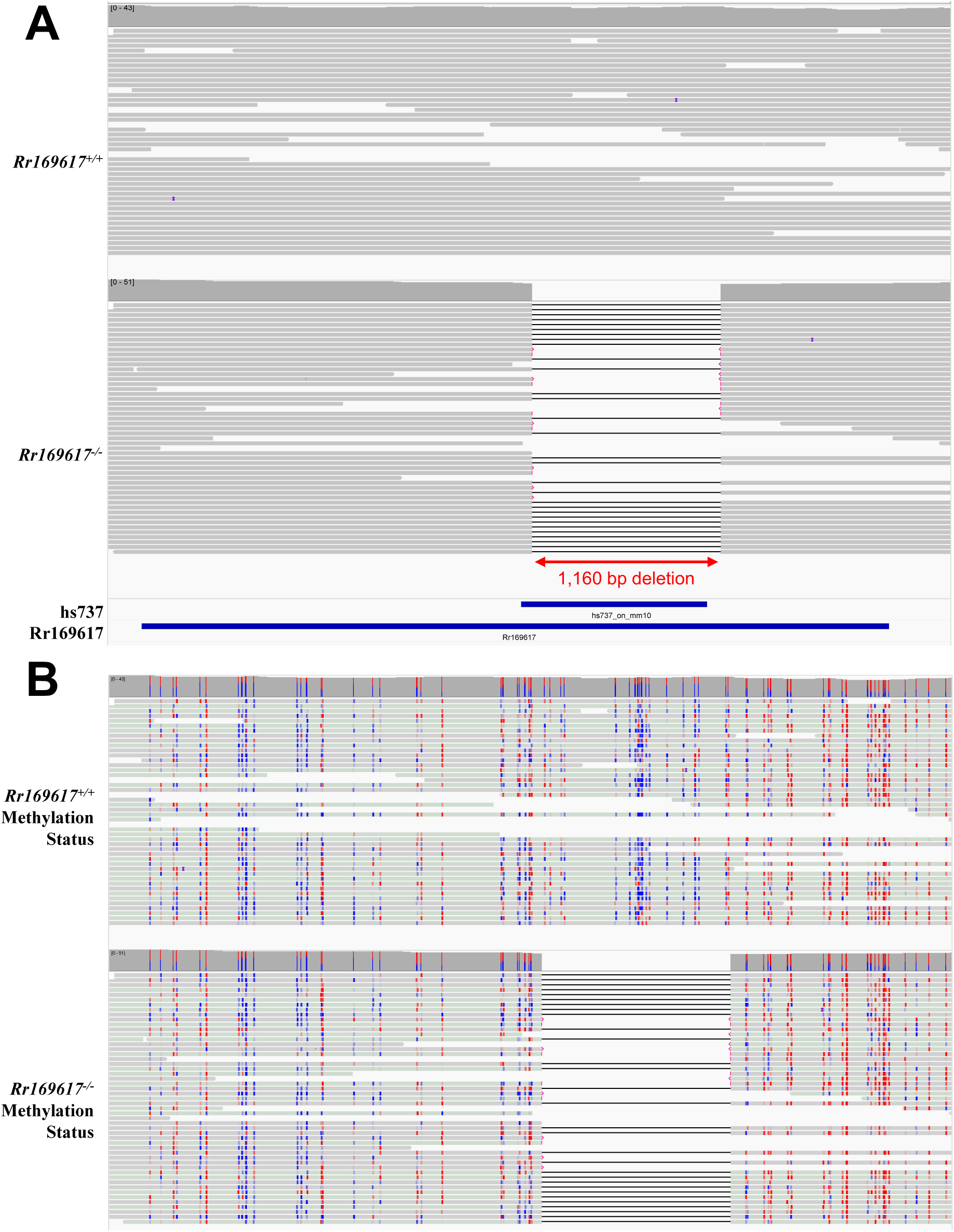
Long-read whole genome sequencing characterization of Cas9 edited mice. A) PacBio HiFi long-read whole-genome sequencing of E12.5 forebrain tissue collected from *Rr169617^+/+^*and *Rr169617^-/-^* mice. All reads in the homozygous deletion mice contain the deletion. None of the reads in the wild type mice contain the deletion. B) Methylation status of CpG sites within the Rr169617 region based on the PacBio whole-genome sequencing data. Note, any of the bases within the portion of Rr169617 containing the sequence orthologous to hs737 are unmethylated as shown in blue. Red = methylated CpG. Blue = unmethylated CpG. For both A and B, the region shown is chr7:136,080,610-136,085,789 (GRCm38/mm10).

Based upon the results of long-read sequencing, we developed a PCR-based genotyping assay that could discriminate between wild type (*Rr169617^+/+^*), heterozygote (*Rr169617^+/-^*), and homozygous deletion (*Rr169617^-/-^*) mice (**Supplementary Figure S2A**) and showed that it matched exactly to the results of whole-genome sequencing (**Supplementary Figure S2B**).

### Underrepresentation of the Deletion Allele

Genotype distributions were analyzed in 278 mice derived from intercrosses between heterozygous animals (*Rr169617^+/-^* × *Rr169617^+/-^*, **Supplementary Figure S3**). From these crosses, we observed 99 (35.6%) wild-type animals *Rr169617^+/+^*, 121 (43.5%) heterozygotes *Rr169617^+/-^*, and 58 (20.9%) homozygotes *Rr169617^-/-^*. This distribution significantly deviates from expected Mendelian frequencies (Chi-Square Test, p = 0.02), demonstrating an underrepresentation of the deletion allele (Binomial test, p = 5.8 × 10^-4^, based on an expected deletion allele frequency of 50%). This showed a confining of homozygous viability but at a reduced level with no significant differences between sexes.

### Consequence of Rr169617 deletion on Ebf3 expression

We hypothesized that deletion in Rr169617 would affect the expression of *Ebf3* based upon the supporting 3D interaction data suggesting that it functions as an enhancer of *Ebf3*. To determine the impact of the deletion on *Ebf3* expression, we collected >10 mouse forebrains, of each genotype, at E12.5 and performed a series of 5 independent qRT-PCR experiments. Consistently, the heterozygous deletion line reduced *Ebf3* expression by ∼10% (**Figure 3**) compared to wild type, and the homozygous deletion line showed a ∼20% reduction in *Ebf3* (**Figure 3**). For significance estimates, we calculated the sample size necessary to detect 80% power for a 20% reduction in expression and found that we need a minimum of 38 samples of each genotype, for 90% power 44 samples of each genotype, and for 100% power 68 samples of each genotype.

**Figure 3:**
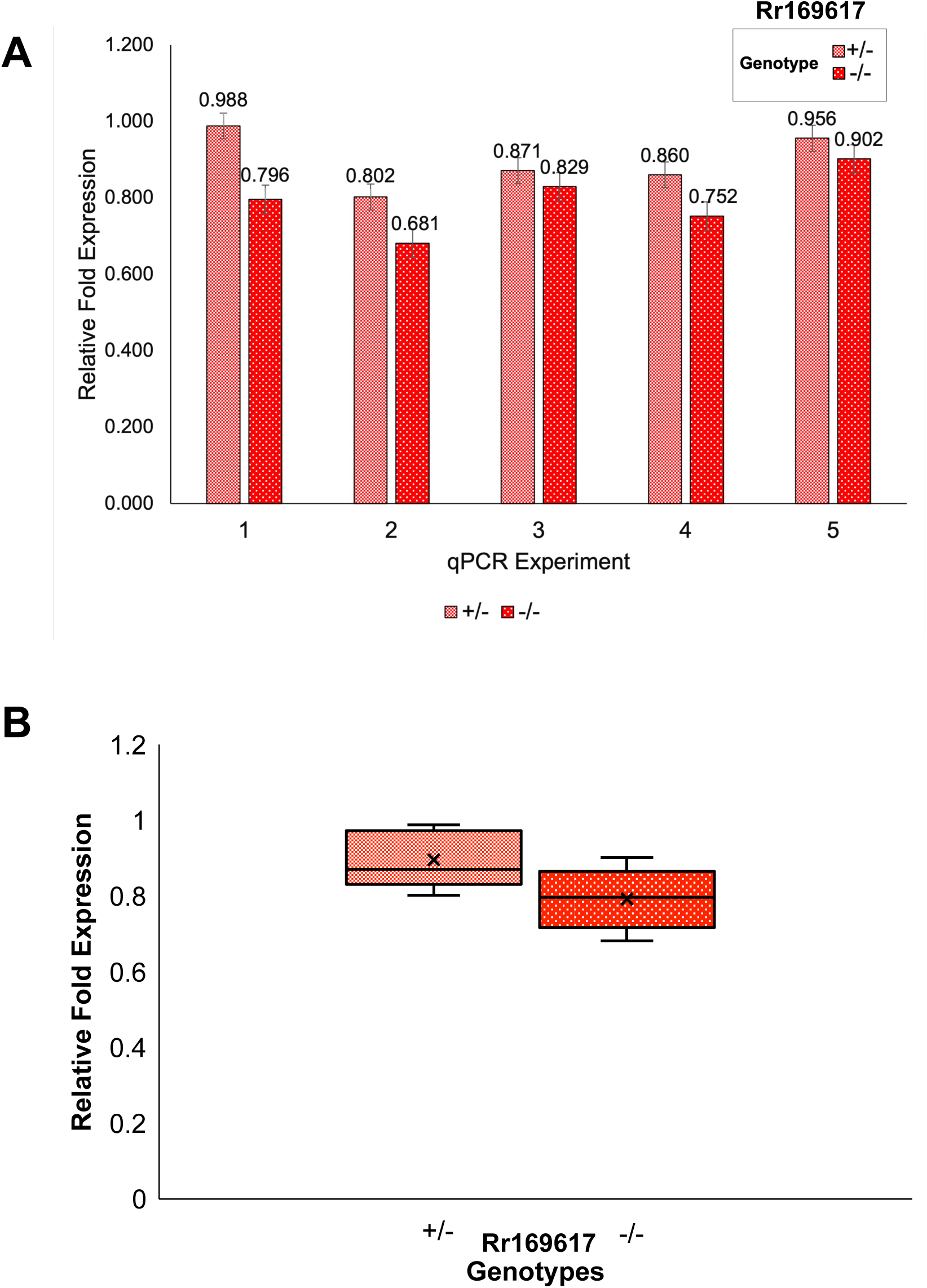
qRT-PCR analysis for *Ebf3* expression in E12.5 forebrain. A) Results of five independent qRT-PCR for *Ebf3* expression. Samples for each genotype were as follows: *Rr169617^+/+^*(n=12), *Rr169617^+/-^* (n=14), and *Rr169617^-/-^*(n=10). B) Relative fold expression aggregating data across all five independent qPCR experiments. For both A and B, relative fold expression is in comparison to the *Rr169617^+/+^* results.

Based on our power estimates, we knew we would not be powered to see a significant difference in *Ebf3* expression in a standard RNAseq experiment because it would be cost prohibitive to sequence a minimum of 76 mice (38 wild type, 38 homozygous deletion, $737 per sample [total of $56,012] just for sequencing). This is an important note for researchers focused on effects of variation in noncoding regions. Therefore, we proceeded to look for downstream (of *Ebf3*) expression changes of much higher effect using a high coverage (∼200 million read pairs) RNAseq experiment on three animals of each genotype in the forebrain, midbrain, and hindbrain, respectively. Through these analyses (**Figure 4, Supplementary Figure S4**), we found there was at least one dysregulated gene in each brain region. In the forebrain (**Supplementary Table 1**), there were 39 genes that were significantly upregulated (17 were protein-coding genes: *Lbhd1*, *Slc4a1*, *Hist1h1e*, *Adra2b*, *Lars2*, *Trim10*, *Nhej1*, *Csf2rb*, *Ncf4*, *Slc25a37*, *Prr15l*, *Acp5*, *Hist1h2bk*, *Hist1h3i*, *Mylk3*, *Cited4*, *Pdzk1ip1*) and 45 genes that were significantly downregulated (11 were protein-coding genes: *Ntng1*, *Ndrg2*, *Hist1h2ao*, *Nox1*, *Neurod6*, *Cst6*, *Cdh12*, *Chd5*, *Htra1*, *Prdm8*, *Glra2*). There were also 14 genes that were significant but did not meet the fold change threshold (11 were protein-coding genes: *Robo2*, *Cachd1*, *B4galt5*, *Bhlhe22*, *Kel*, *Hbb-y*, *Hsd3b6*, *Csf2ra*, *Cabp1*, *Asrgl1*, *Hspa8*). In the midbrain (**Supplementary Table S2**), there was only 1 gene that was significantly downregulated (pseudogene *Pisd-ps1*). In the hindbrain (**Supplementary Table S3**), there were 5 genes that were significantly upregulated (1 protein-coding: *mt-Atp6*) and 8 genes that were significantly downregulated (4 were protein-coding genes: *Mroh7*, *Col7a1*, *Ndor1*, *Trpc2*). There were also 10 genes that were significant but did not meet the fold change threshold (6 were protein-coding: *Pisd*, *Manea*, *mt-Nd2*, *Ldb2*, *Aif1l*, *Aldh16a1*).

**Figure 4:**
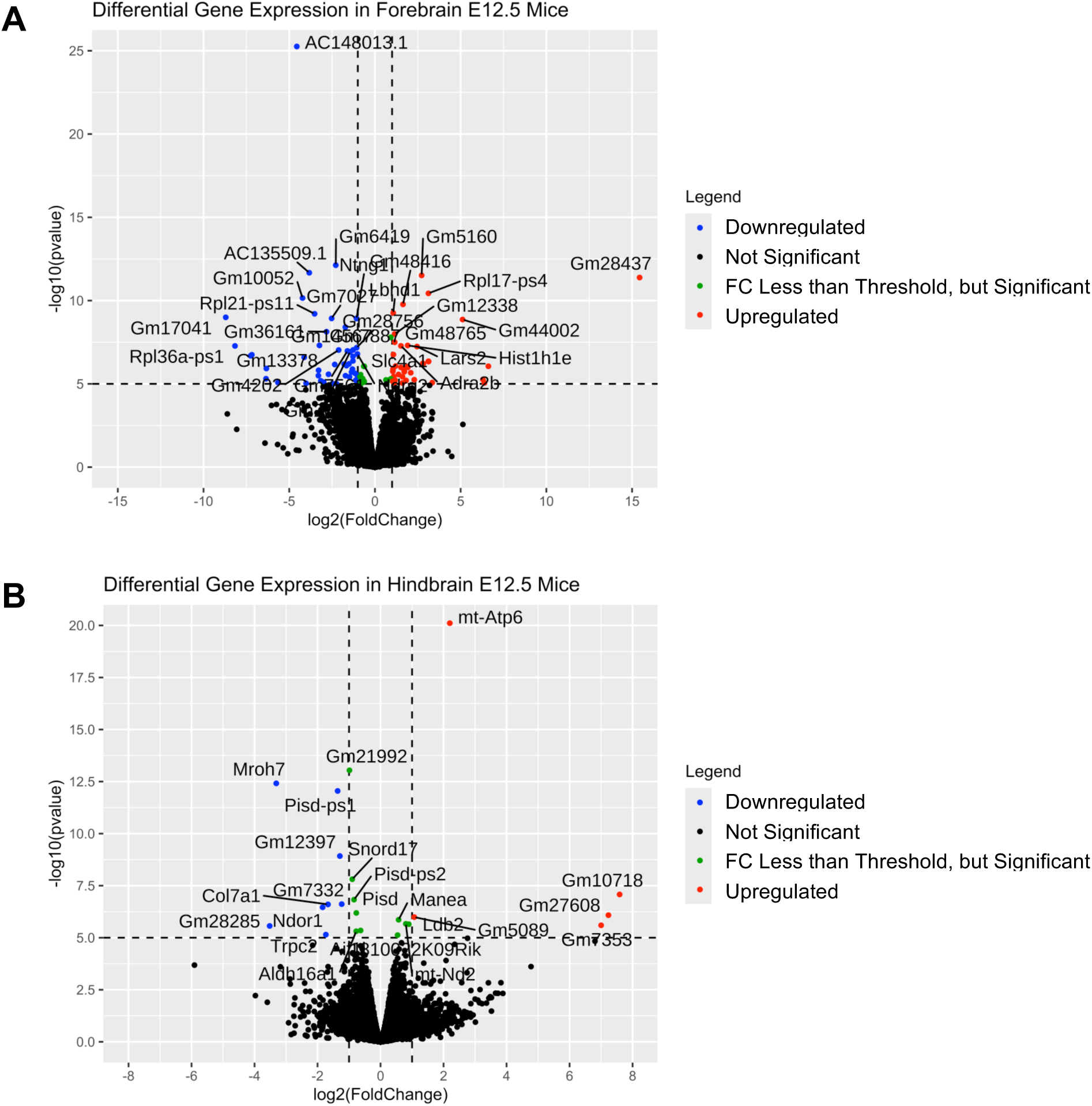
Volcano plots of RNA-Seq data from E12.5 mice. Volcano plots from A) forebrain and B) hindbrain mice collected at E12.5 are shown. Each figure compares a *Rr169617^-/-^*vs. *Rr169617^+/+^*. The fold change cutoff was Log2>=|1|, and a p-value cutoff of >=10e-6 was used, denoted by the dashed black lines. Red dots are significantly up-regulated genes, blue dots are significantly down-regulated genes, and green dots are genes that met statistical significance but did not meet the fold change threshold.

### Phenotyping Analysis of Enhancer Deletion Mice

Given the reproducible reduction in expression of *Ebf3* observed in our enhancer deletion mice, we next performed phenotyping tests to assess potential behavioral differences focusing on hole board (HB), light-dark transition (LD) and open field (OF). These tests allowed us to assess traits related to anxiety, exploration and mobility. To increase power and show generalizability, the percentage time of the mouse was mobile (locomotion) was compared as repeated measures over the three behavioral assays (OF, HB, L/D). In all cases the *Rr169617^-/-^* showed a difference between the sexes for time mobile with males less active than females (overall p = 0.005: by assay, OF 0.045; LD 0.001; HB 0.041)). Wildtype mice showed no significant sexual dimorphism (p = 0.34; **Figure 5A**). Similarly, the two assays also provide an assessment of speed of travel (OF, HB) and showed sexual dimorphism only for *Rr169617^-/-^* with males slower than females (overall p = 0.003:, by assay OF 0.01; HB 0.002), wildtype mice showed no difference between the sexes.

**Figure 5:**
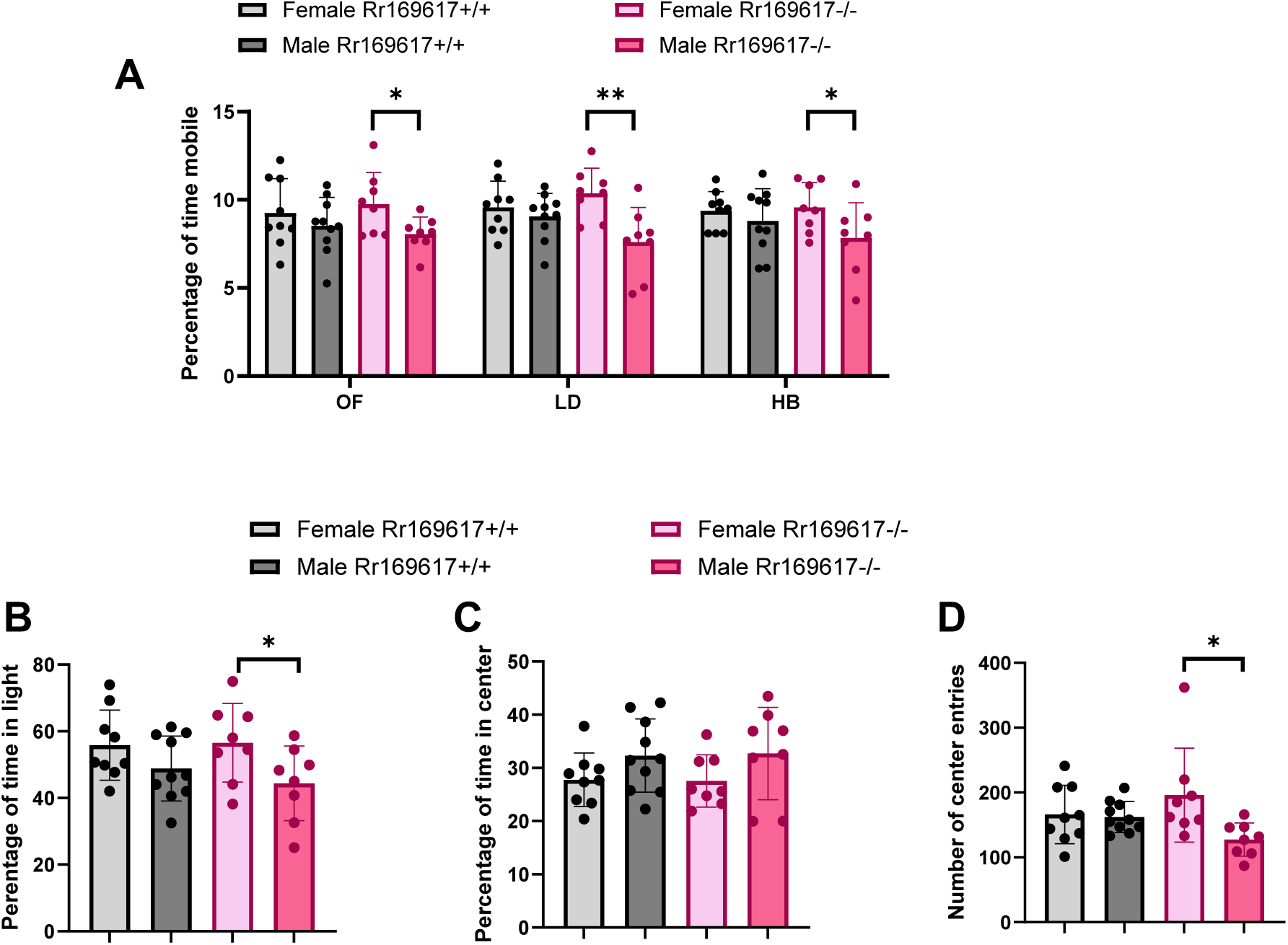
Behavioral phenotypes observed in Rr169617 deletion mice. (A) Summary of mouse mobility as measured by three independent assays: open field (OF), light-dark (LD), and hole board (HB). In all assays, males homozygous for deletion of Rr169617 were less active than homozygous females. (B,C,D) Summary of additional parameters measured in the light-dark assay.(B) and open field (C,D) assays. Within strain, sex-specific differences were observed when comparing homozygous deletion males versus females. (B) Homozygous males spent less time in the light and (D) showed fewer center entries; (C) however, no significant difference was observed for the time spent in center. Asterisk (*) indicates statistical significance (p < 0.05) between groups.

In addition to mobility, the light/dark and open field assays provide a measure of potential anxiety with anxious mice predicted to spend less time in the center of the open field and less time in the light portion of the LD box. The LD assay revealed significant sexual dimorphism effects for *Rr169617^-/-^* not seen in *Rr169617^+/+^* mice. The *Rr169617^-/-^* males spent less time in the light than females indicating a possible higher level of anxiety (p = 0.03, **Figure 5B**). However, OF did not confirm this result as no differences were found for the strains for the time in center (p=0.13, **Figure 5C**). Both the *a priori* and Phenstat analyses showed a sex difference for the number of center entries in OF. The *a priori* analysis determined that the male *Rr169617^-/-^* mice showed fewer entries to center than females (p=0.005, **Figure 5D**) which was not seen for the wildtype mice (p =0.855). The PhenStat analysis revealed sexual dimorphism such that male *Rr169617^-/-^* mice showed significantly fewer entries compared to wildtype males (percent effect change = -21.35% : p=0.04) and the *Rr169617^-/-^* females showed significantly more entries compared to wildtype females (percent effect change = 18.33%). The number of entries into the center is a weak indicator of anxiety as it is confounded by a clear difference in the amount of movement between the male and female *Rr169617^-/-^* mice.

The behavioral assays described above were performed as part of a broad-based phenotyping pipeline implemented by the International Mouse Phenotyping Consortium ^46^. All mice in this study were also assessed for a variety of other physiological measures that includes clinical blood chemistry, body composition via DEXA, and gross morphological assessment. Notably, we detected difference in body composition parameters that showed an overall gene effect as *Rr169617^-/-^* mice had a lower proportion of fat than wildtype mice and conversely an increased lean proportion (e.g. fat(g)/bodyweight (g) p = 0.013, lean mass (g)/bodyweight (g) p =0.006). Measures of the bone parameters revealed that the male *Rr169617^-/-^* mice have larger, denser bones than females (e.g. Bone Mineral Density p=0.001, Bone Area p = 0.003) but that the wildtype mice showed no sexual dimorphism for these bone parameters. As expected, parameters that assessed body size showed a sex difference for both *Rr169617^-/-^* and *Rr169617^+/+^* mice (e.g. bodyweight p<0.001; body length p<0.001) with males larger than females. While we analyzed all phenotyping tests performed using the PhenStats packaged, it revealed very few significant genotype-specific effects (see Supplemental Tables S6 to S20). The only significant gene effects found were for the DEXA parameters of the relative proportions of fat and lean mass already discussed.

## DISCUSSION

Discovery of genes involved in autism and other neurodevelopmental disorders is occurring rapidly with the utility of several sequencing strategies. Moving beyond the exome, there is an appreciation that noncoding regions of the genome also play an important role. These regions finely tune the expression of genes and are particularly important in brain development. Several areas are being pursued with regard to noncoding regions including statistical testing for enrichment, machine-learning based models, functional characterization at thousands of regions/variants at a time (i.e., Multiplex Assays of Variant Effects), transient transgenic assays, and through precision engineering in model organisms. In this study, we follow up on our previous identification of variants in individuals with neurodevelopmental disorders in the enhancer hs737 that affects the target gene, *EBF3*. *EBF3* is well-established as a syndromic gene and has genome-wide significance for excess variation in individuals with neurodevelopmental disorders. However, the knowledge of the function of this gene and the gene regulatory network (GRN) that it resides within are understudied at this time. Our identification of hs737 provides a foothold into looking at the upstream regulators of EBF3. Further, we observed an excess of deletions of hs737 in individuals with neurodevelopmental disorders consistent with the finding that both EBF3 and its associated regulatory elements are required for normal neural development.

In this study, we pursued the hypothesis that deletion of this element in the highly conserved, orthologous mouse region would provide additional insights into hs737/Rr169617 and *EBF3*/*Ebf3*. First, we found that when the corresponding region in mouse of hs737 is deleted it results in fewer progeny homozygous for the deletion than expected by chance. This suggests that while homozygous deletion mice are viable they are not being born at the rate expected by Mendelian inheritance. Second, mice homozygous for this deletion displayed a relatively small effect on expression of the target gene. However, each enhancer has a different effect on expression, and it is not readily apparent *a priori* what the reduction of a specific enhancer will be in an *in vivo* model. Thus, emphasizing the critical importance of experimentally testing regulatory sequences in the context of a whole animal. The modest reduction in expression can also have consequences for RNAseq experiment design; whereby, exorbitant sample sizes would be necessary to see the expression difference as significant. Therefore, we recommend that smaller sample sizes may be pursued for the RNAseq experiments but that the outcome of these experiments will only reveal the genes with highest changes in expression. We identified genes with high changes in expression in this study including *Lbhd1* and *Ntng1*. Third, we identified sex specific phenotypes related to mobility and anxiety when comparing males versus females homozygous for the enhancer deletion. This is highly relevant to the phenotype of autism and the phenotypes we see in individuals with variation in the hs737 enhancer (males with autism and hypotonia but no intellectual disability) ^33^.

In future studies, it will be important to identify the identity of the upstream transcription factors that bind hs737, determine its activity at the single-cell level, and characterize the other cis-regulatory elements that help orchestrate the precise developmental expression of *Ebf3*. These analyses will provide a framework for investigating the effects of other non-coding variants found within the *Ebf3* regulatory landscape. Here, we focused on the deletion of a single element due to the overwhelming supporting evidence from individuals with neurodevelopmental disorders that harbor deletions in this region. The detailed characterization of this novel mouse model provides insight into the molecular and phenotypic impacts of deleting this enhancer and will be of interest to the broad biomedical research community interested in understanding how changes in the noncoding fraction of the genome affect human health and disease.

## ACKNOWLEDGMENTS

This work was supported by grants from the National Institutes of Health (R00MH117165 to T.N.T., R01MH126933 to T.N.T., UM1OD023222 to S.A.M and J.K.W, and R01HD102534 D.U.G.) and the McDonnell Center for Cellular and Molecular Neurobiology at Washington University in St. Louis. The content is solely the responsibility of the authors and does not necessarily represent the official views of the National Institutes of Health.

## DATA AND SOFTWARE AVAILABILITY STATEMENT

Data is available at NCBI BioProject PRJNA1194105.

## SUPPLEMENTARY FIGURE LEGENDS

**Supplementary Figure S1:**
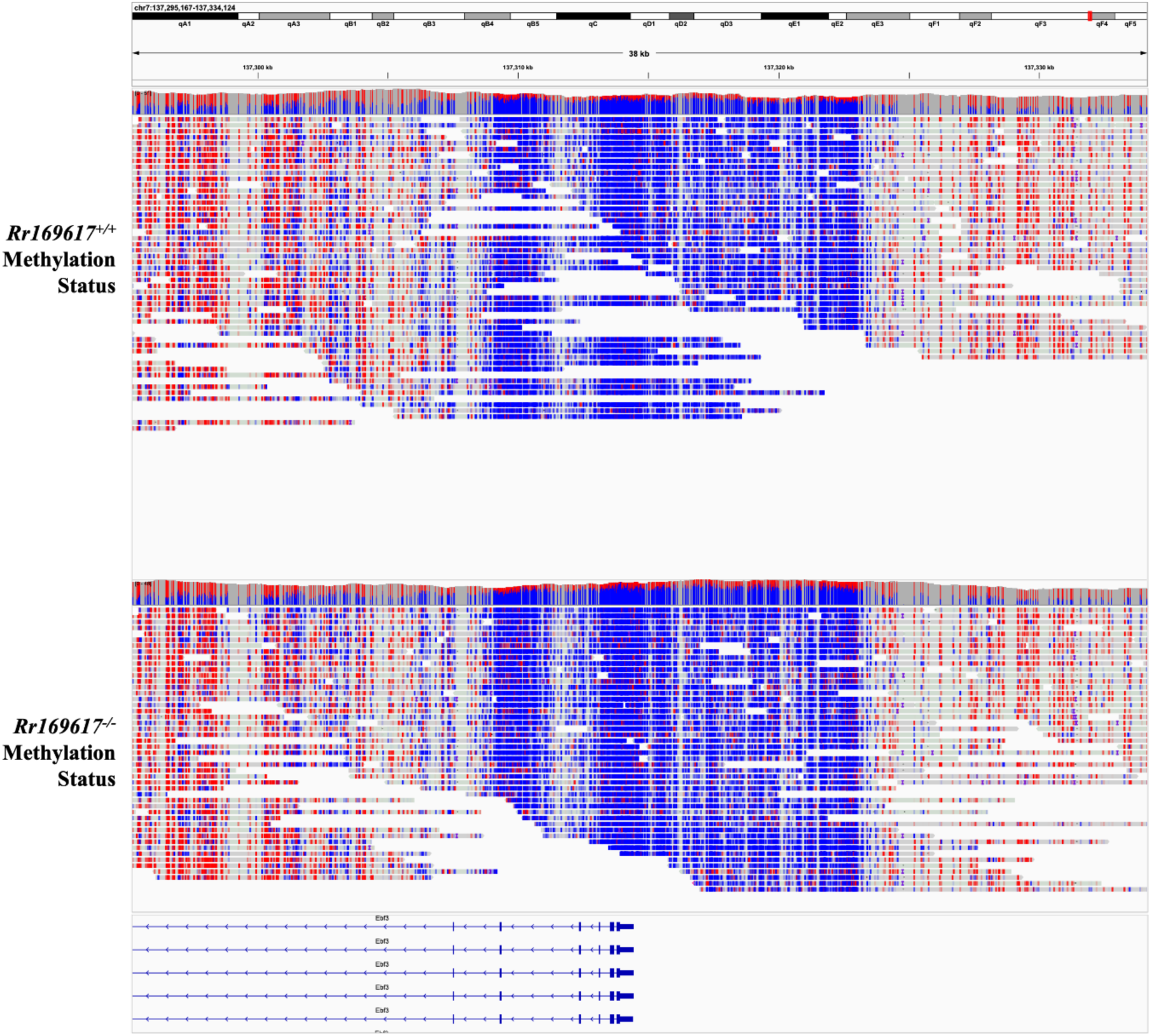
Methylation Status at the *Ebf3* Promoter in *Rr169617^+/+^* and *Rr169617^-/-^*E12.5 forebrains. Shown is the methylation status of CpG sites within the *Ebf3* promoter region based on the PacBio whole-genome sequencing data. The methylation patterns look similar in both. Red = methylated CpG. Blue = unmethylated CpG.

**Supplementary Figure S2:**
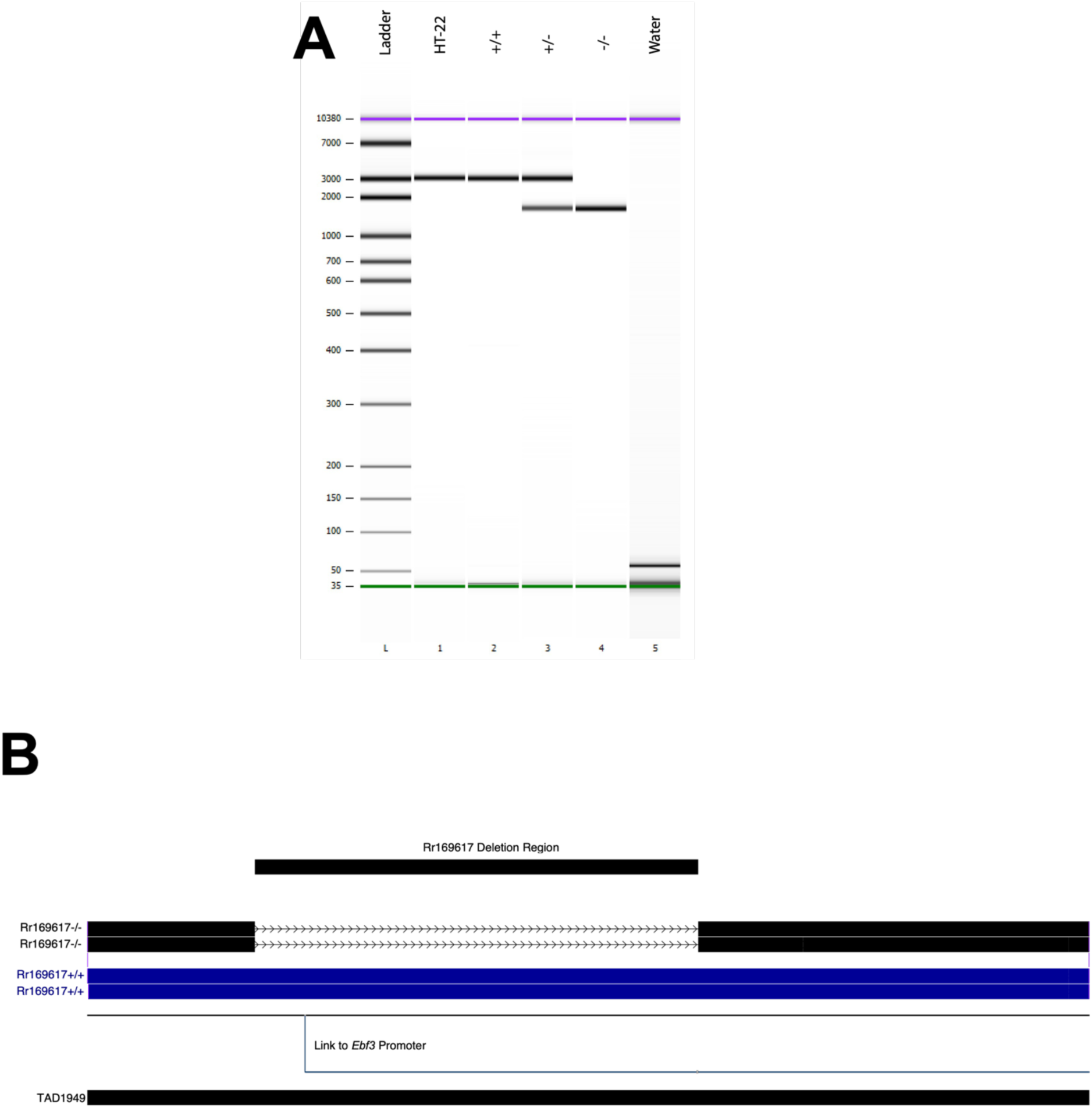
Genotyping assay for 1,160 bp deletion in Rr169617 mice. A) Results of PCR-based assay to genotype for the deletion. B) Sequencing of PCR products confirming they match the *Rr169617^+/+^* and *Rr169617^-/-^* expected results. Shown are two replicates of each.

**Supplementary Figure S3:**
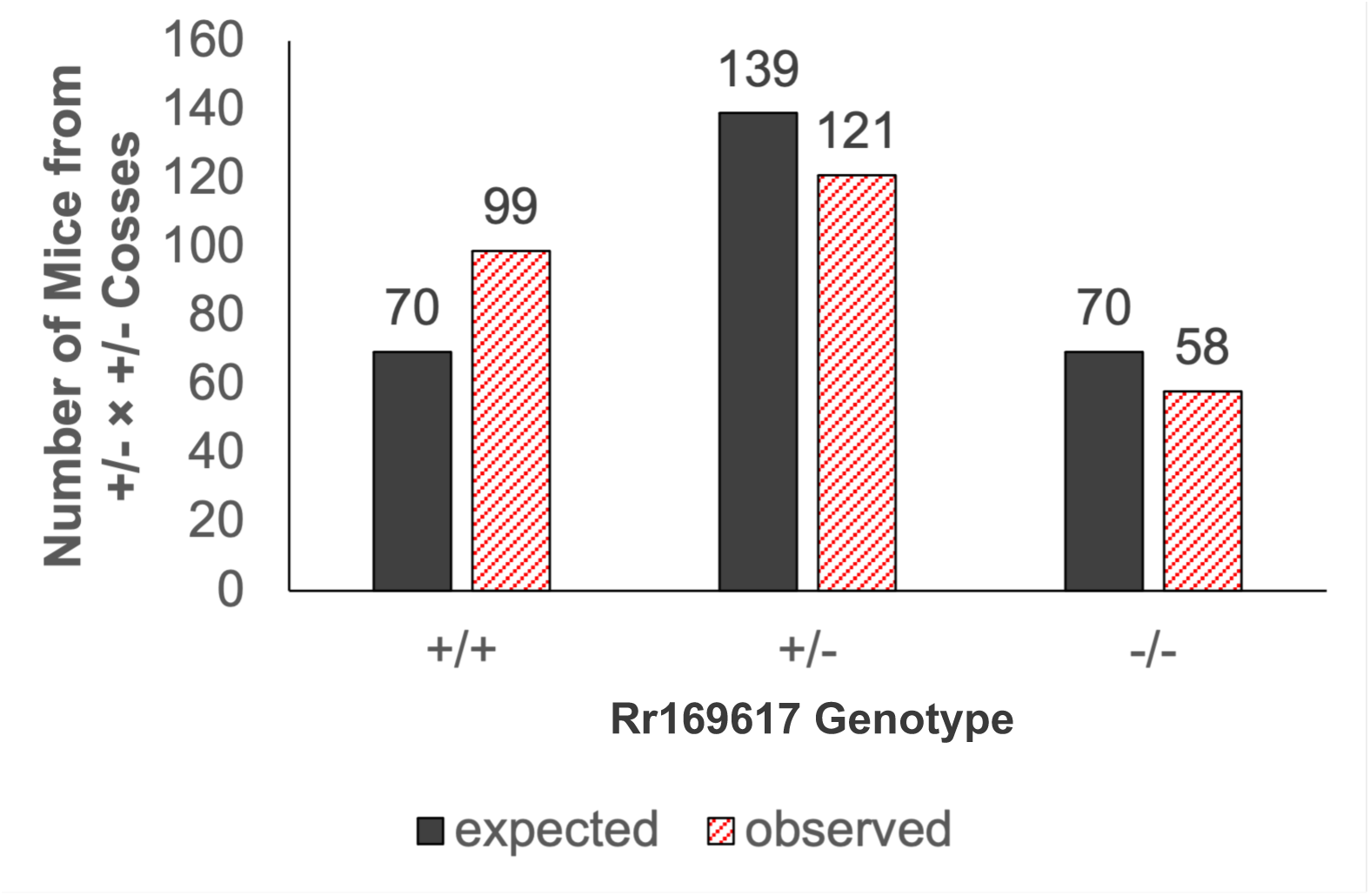
Genotypes of 278 Mice from *Rr169617^+/-^* × *Rr169617^+/-^* Crosses. Shown in black are the expected counts (based on Mendelian inheritance) and in red are the actual observed counts from heterozygote crosses. This distribution significantly deviates from expected Mendelian frequencies (Chi-Square Test, p = 0.02).

**Supplementary Figure S4:**
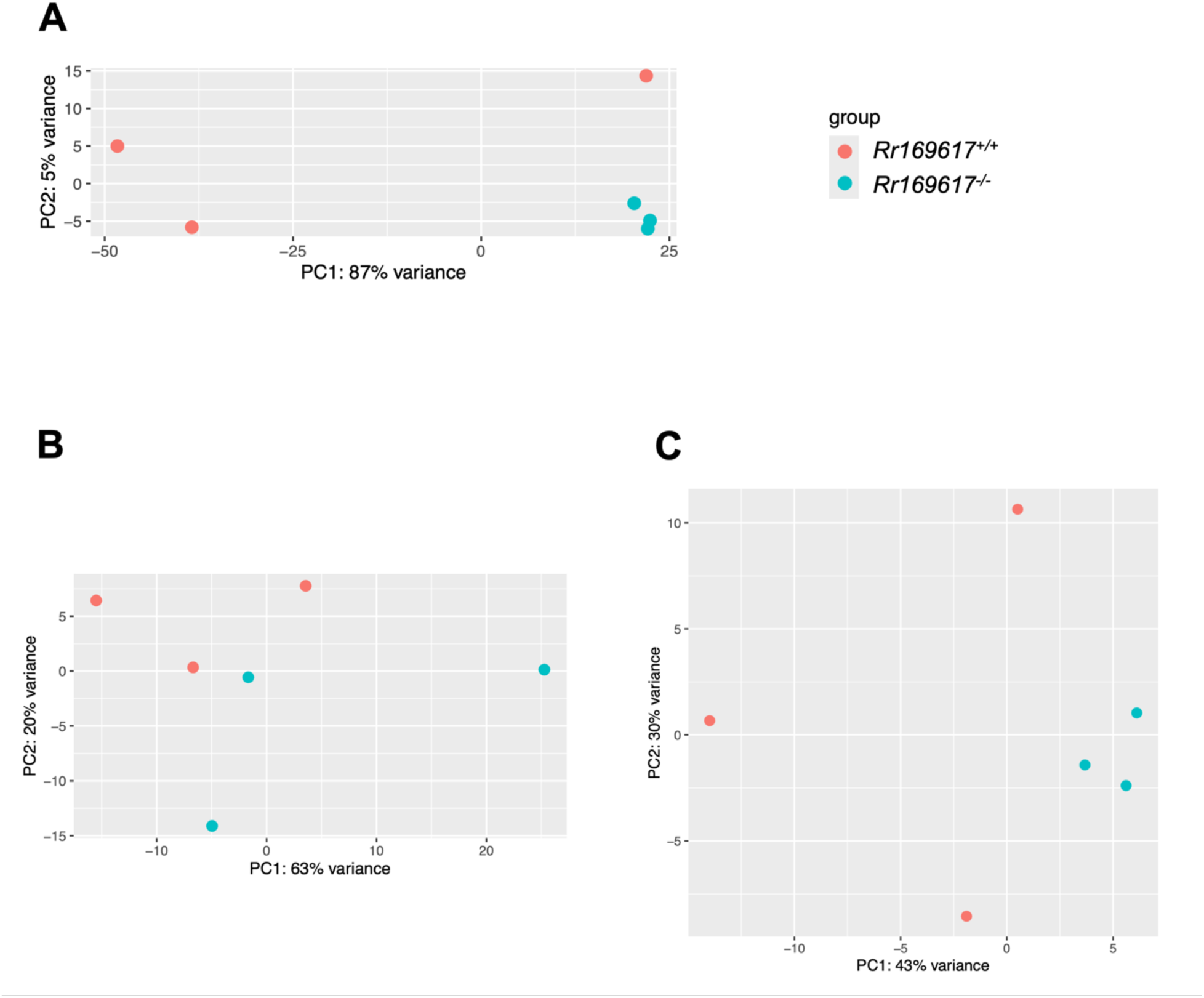
PCA plots of RNA-seq data from E12.5 *Rr169617^-/-^* and *Rr169617^+/+^* mice. PCA plots showing the clustering of the wild-type and homozygous deletion samples in A) forebrain, B) rRNA-depleted midbrain, and C) rRNA-depleted hindbrain mice collected at E12.5.

## SUPPLEMENTARY TABLE LEGENDS

**Supplementary Table S1.** E12.5 Forebrain RNAseq Results

**Supplementary Table S2.** E12.5 Midbrain RNAseq Results

**Supplementary Table S3.** E12.5 Hindbrain RNAseq Results

**Supplementary Table S4.** Mouse Details for the KOMP Phenotyping

**Supplementary Table S5.** KOMP Phenotyping Pipeline

**Supplementary Table S6.** KOMP Phenotyping Standalone Body Weight

**Supplementary Table S7.** KOMP Phenotyping Open Field

**Supplementary Table S8.** KOMP Phenotyping SHIRPA Dysmorphology

**Supplementary Table S9.** KOMP Phenotyping Grip Strength

**Supplementary Table S10.** KOMP Phenotyping Light Dark Transition

**Supplementary Table S11.** KOMP Phenotyping Holeboard

**Supplementary Table S12.** KOMP Phenotyping Acoustic Startle PPI

**Supplementary Table S13.** KOMP Phenotyping Electrocardiography

**Supplementary Table S14.** KOMP Phenotyping Glucose Tolerance

**Supplementary Table S15.** KOMP Phenotyping Body Composition

**Supplementary Table S16.** KOMP Phenotyping Eye Morphology

**Supplementary Table S17.** KOMP Phenotyping Auditory Brainstem Response

**Supplementary Table S18.** KOMP Phenotyping Hematology

**Supplementary Table S19.** KOMP Phenotyping Clinical Blood Chemistry

**Supplementary Table S20.** KOMP Phenotyping Heart Weight

## REFERENCES

1. Bai, D., Yip, B.H.K., Windham, G.C., Sourander, A., Francis, R., Yoffe, R., Glasson, E., Mahjani, B., Suominen, A., Leonard, H., et al. (2019). Association of Genetic and Environmental Factors With Autism in a 5-Country Cohort. JAMA Psychiatry.

2. Sandin, S., Lichtenstein, P., Kuja-Halkola, R., Hultman, C., Larsson, H., and Reichenberg, A. (2017). The heritability of autism spectrum disorder. Jama 318, 1182–1184.

3. De Rubeis, S., He, X., Goldberg, A.P., Poultney, C.S., Samocha, K., Cicek, A.E., Kou, Y., Liu, L., Fromer, M., Walker, S., et al. (2014). Synaptic, transcriptional and chromatin genes disrupted in autism. Nature 515, 209–215.

4. Dong, S., Walker, M.F., Carriero, N.J., DiCola, M., Willsey, A.J., Ye, A.Y., Waqar, Z., Gonzalez, L.E., Overton, J.D., Frahm, S., et al. (2014). De novo insertions and deletions of predominantly paternal origin are associated with autism spectrum disorder. Cell reports 9, 16–23.

5. Iossifov, I., O’Roak, B.J., Sanders, S.J., Ronemus, M., Krumm, N., Levy, D., Stessman, H.A., Witherspoon, K.T., Vives, L., Patterson, K.E., et al. (2014). The contribution of de novo coding mutations to autism spectrum disorder. Nature.

6. Sanders, S.J., Murtha, M.T., Gupta, A.R., Murdoch, J.D., Raubeson, M.J., Willsey, A.J., Ercan-Sencicek, A.G., DiLullo, N.M., Parikshak, N.N., Stein, J.L., et al. (2012). De novo mutations revealed by whole-exome sequencing are strongly associated with autism. Nature 485, 237–241.

7. Iossifov, I., Ronemus, M., Levy, D., Wang, Z., Hakker, I., Rosenbaum, J., Yamrom, B., Lee, Y.H., Narzisi, G., Leotta, A., et al. (2012). De novo gene disruptions in children on the autistic spectrum. Neuron 74, 285–299.

8. Krumm, N., Turner, T.N., Baker, C., Vives, L., Mohajeri, K., Witherspoon, K., Raja, A., Coe, B.P., Stessman, H.A., He, Z.X., et al. (2015). Excess of rare, inherited truncating mutations in autism. Nature genetics 47, 582–588.

9. O’Roak, B.J., Deriziotis, P., Lee, C., Vives, L., Schwartz, J.J., Girirajan, S., Karakoc, E., Mackenzie, A.P., Ng, S.B., Baker, C., et al. (2011). Exome sequencing in sporadic autism spectrum disorders identifies severe de novo mutations. Nature genetics 43, 585–589.

10. O’Roak, B.J., Stessman, H.A., Boyle, E.A., Witherspoon, K.T., Martin, B., Lee, C., Vives, L., Baker, C., Hiatt, J.B., Nickerson, D.A., et al. (2014). Recurrent de novo mutations implicate novel genes underlying simplex autism risk. Nature communications 5, 5595.

11. O’Roak, B.J., Vives, L., Fu, W., Egertson, J.D., Stanaway, I.B., Phelps, I.G., Carvill, G., Kumar, A., Lee, C., Ankenman, K., et al. (2012). Multiplex targeted sequencing identifies recurrently mutated genes in autism spectrum disorders. Science (New York, NY) 338, 1619–1622.

12. O’Roak, B.J., Vives, L., Girirajan, S., Karakoc, E., Krumm, N., Coe, B.P., Levy, R., Ko, A., Lee, C., Smith, J.D., et al. (2012). Sporadic autism exomes reveal a highly interconnected protein network of de novo mutations. Nature 485, 246–250.

13. De Rubeis, S., and Buxbaum, J.D. (2015). Genetics and genomics of autism spectrum disorder: embracing complexity. Hum Mol Genet 24, R24–31.

14. Coe, B.P., Witherspoon, K., Rosenfeld, J.A., van Bon, B.W., Vulto-van Silfhout, A.T., Bosco, P., Friend, K.L., Baker, C., Buono, S., Vissers, L.E., et al. (2014). Refining analyses of copy number variation identifies specific genes associated with developmental delay. Nature genetics 46, 1063–1071.

15. Cooper, G.M., Coe, B.P., Girirajan, S., Rosenfeld, J.A., Vu, T.H., Baker, C., Williams, C., Stalker, H., Hamid, R., Hannig, V., et al. (2011). A copy number variation morbidity map of developmental delay. Nature genetics 43, 838–846.

16. Girirajan, S., Dennis, M.Y., Baker, C., Malig, M., Coe, B.P., Campbell, C.D., Mark, K., Vu, T.H., Alkan, C., Cheng, Z., et al. (2013). Refinement and discovery of new hotspots of copy-number variation associated with autism spectrum disorder. American journal of human genetics 92, 221–237.

17. Krumm, N., O’Roak, B.J., Karakoc, E., Mohajeri, K., Nelson, B., Vives, L., Jacquemont, S., Munson, J., Bernier, R., and Eichler, E.E. (2013). Transmission disequilibrium of small CNVs in simplex autism. American journal of human genetics 93, 595–606.

18. Sanders, S.J., Ercan-Sencicek, A.G., Hus, V., Luo, R., Murtha, M.T., Moreno-De-Luca, D., Chu, S.H., Moreau, M.P., Gupta, A.R., Thomson, S.A., et al. (2011). Multiple recurrent de novo CNVs, including duplications of the 7q11.23 Williams syndrome region, are strongly associated with autism. Neuron 70, 863–885.

19. Sanders, S.J., He, X., Willsey, A.J., Ercan-Sencicek, A.G., Samocha, K.E., Cicek, A.E., Murtha, M.T., Bal, V.H., Bishop, S.L., Dong, S., et al. (2015). Insights into Autism Spectrum Disorder Genomic Architecture and Biology from 71 Risk Loci. Neuron 87, 1215–1233.

20. Levy, D., Ronemus, M., Yamrom, B., Lee, Y.H., Leotta, A., Kendall, J., Marks, S., Lakshmi, B., Pai, D., Ye, K., et al. (2011). Rare de novo and transmitted copy-number variation in autistic spectrum disorders. Neuron 70, 886–897.

21. Marshall, C.R., Noor, A., Vincent, J.B., Lionel, A.C., Feuk, L., Skaug, J., Shago, M., Moessner, R., Pinto, D., Ren, Y., et al. (2008). Structural variation of chromosomes in autism spectrum disorder. American journal of human genetics 82, 477–488.

22. Pinto, D., Pagnamenta, A.T., Klei, L., Anney, R., Merico, D., Regan, R., Conroy, J., Magalhaes, T.R., Correia, C., Abrahams, B.S., et al. (2010). Functional impact of global rare copy number variation in autism spectrum disorders. Nature 466, 368–372.

23. Weiner, D.J., Wigdor, E.M., Ripke, S., Walters, R.K., Kosmicki, J.A., Grove, J., Samocha, K.E., Goldstein, J.I., Okbay, A., Bybjerg-Grauholm, J., et al. (2017). Polygenic transmission disequilibrium confirms that common and rare variation act additively to create risk for autism spectrum disorders. Nature genetics 49, 978–985.

24. Turner, T.N., Coe, B.P., Dickel, D.E., Hoekzema, K., Nelson, B.J., Zody, M.C., Kronenberg, Z.N., Hormozdiari, F., Raja, A., Pennacchio, L.A., et al. (2017). Genomic Patterns of De Novo Mutation in Simplex Autism. Cell 171, 710–722.e712.

25. Turner, T.N., Hormozdiari, F., Duyzend, M.H., McClymont, S.A., Hook, P.W., Iossifov, I., Raja, A., Baker, C., Hoekzema, K., Stessman, H.A., et al. (2016). Genome Sequencing of Autism-Affected Families Reveals Disruption of Putative Noncoding Regulatory DNA. American journal of human genetics 98, 58–74.

26. An, J.Y., Lin, K., Zhu, L., Werling, D.M., Dong, S., Brand, H., Wang, H.Z., Zhao, X., Schwartz, G.B., Collins, R.L., et al. (2018). Genome-wide de novo risk score implicates promoter variation in autism spectrum disorder. Science (New York, NY) 362.

27. Werling, D.M., Brand, H., An, J.Y., Stone, M.R., Zhu, L., Glessner, J.T., Collins, R.L., Dong, S., Layer, R.M., Markenscoff-Papadimitriou, E., et al. (2018). An analytical framework for whole-genome sequence association studies and its implications for autism spectrum disorder. Nature genetics 50, 727–736.

28. Brandler, W.M., Antaki, D., Gujral, M., Kleiber, M.L., Whitney, J., Maile, M.S., Hong, O., Chapman, T.R., Tan, S., Tandon, P., et al. (2018). Paternally inherited cis-regulatory structural variants are associated with autism. Science (New York, NY) 360, 327–331.

29. Brandler, W.M., Antaki, D., Gujral, M., Noor, A., Rosanio, G., Chapman, T.R., Barrera, D.J., Lin, G.N., Malhotra, D., Watts, A.C., et al. (2016). Frequency and Complexity of De Novo Structural Mutation in Autism. American journal of human genetics 98, 667–679.

30. Padhi, E.M., Hayeck, T.J., Mannion, B., Chatterjee, S., Byrska-Bishop, M., Musunuri, R., Narzisi, G., Abhyankar, A., Cheng, Z., Hunter, R.D., et al. (2020). De Novo Mutation in an Enhancer of EBF3 in simplex autism. bioRxiv, 2020.2008.2028.270751.

31. Markenscoff-Papadimitriou, E., Whalen, S., Przytycki, P., Thomas, R., Binyameen, F., Nowakowski, T.J., Kriegstein, A.R., Sanders, S.J., State, M.W., Pollard, K.S., et al. (2020). A Chromatin Accessibility Atlas of the Developing Human Telencephalon. Cell 182, 754–769.e718.

32. Zhou, J., Park, C.Y., Theesfeld, C.L., Wong, A.K., Yuan, Y., Scheckel, C., Fak, J.J., Funk, J., Yao, K., Tajima, Y., et al. (2019). Whole-genome deep-learning analysis identifies contribution of noncoding mutations to autism risk. Nature genetics 51, 973–980.

33. Padhi, E.M., Hayeck, T.J., Cheng, Z., Chatterjee, S., Mannion, B.J., Byrska-Bishop, M., Willems, M., Pinson, L., Redon, S., Benech, C., et al. (2021). Coding and noncoding variants in EBF3 are involved in HADDS and simplex autism. Hum Genomics 15, 44.

34. Chao, H.T., Davids, M., Burke, E., Pappas, J.G., Rosenfeld, J.A., McCarty, A.J., Davis, T., Wolfe, L., Toro, C., Tifft, C., et al. (2017). A Syndromic Neurodevelopmental Disorder Caused by De Novo Variants in EBF3. American journal of human genetics 100, 128–137.

35. Harms, F.L., Girisha, K.M., Hardigan, A.A., Kortüm, F., Shukla, A., Alawi, M., Dalal, A., Brady, L., Tarnopolsky, M., Bird, L.M., et al. (2017). Mutations in EBF3 Disturb Transcriptional Profiles and Cause Intellectual Disability, Ataxia, and Facial Dysmorphism. American journal of human genetics 100, 117–127.

36. Sleven, H., Welsh, S.J., Yu, J., Churchill, M.E.A., Wright, C.F., Henderson, A., Horvath, R., Rankin, J., Vogt, J., Magee, A., et al. (2017). De Novo Mutations in EBF3 Cause a Neurodevelopmental Syndrome. American journal of human genetics 100, 138–150.

37. Ignatius, E., Puosi, R., Palomäki, M., Forsbom, N., Pohjanpelto, M., Alitalo, T., Anttonen, A.K., Avela, K., Haataja, L., Carroll, C.J., et al. (2022). Duplication/triplication mosaicism of EBF3 and expansion of the EBF3 neurodevelopmental disorder phenotype. Eur J Paediatr Neurol 37, 1–7.

38. Liberg, D., Sigvardsson, M., and Akerblad, P. (2002). The EBF/Olf/Collier family of transcription factors: regulators of differentiation in cells originating from all three embryonal germ layers. Molecular and cellular biology 22, 8389–8397.

39. Chen, Z., Snetkova, V., Bower, G., Jacinto, S., Clock, B., Dizehchi, A., Barozzi, I., Mannion, B.J., Alcaina-Caro, A., Lopez-Rios, J., et al. (2024). Increased enhancer-promoter interactions during developmental enhancer activation in mammals. Nature genetics 56, 675–685.

40. Pennacchio, L.A., Ahituv, N., Moses, A.M., Prabhakar, S., Nobrega, M.A., Shoukry, M., Minovitsky, S., Dubchak, I., Holt, A., Lewis, K.D., et al. (2006). In vivo enhancer analysis of human conserved non-coding sequences. Nature 444, 499–502.

41. Love, M.I., Huber, W., and Anders, S. (2014). Moderated estimation of fold change and dispersion for RNA-seq data with DESeq2. Genome biology 15, 550.

42. Dickinson, M.E., Flenniken, A.M., Ji, X., Teboul, L., Wong, M.D., White, J.K., Meehan, T.F., Weninger, W.J., Westerberg, H., Adissu, H., et al. (2016). High-throughput discovery of novel developmental phenotypes. Nature 537, 508–514.

43. Kurbatova, N., Mason, J.C., Morgan, H., Meehan, T.F., and Karp, N.A. (2015). PhenStat: A Tool Kit for Standardized Analysis of High Throughput Phenotypic Data. PLoS One 10, e0131274.

44. Dixon, J.R., Selvaraj, S., Yue, F., Kim, A., Li, Y., Shen, Y., Hu, M., Liu, J.S., and Ren, B. (2012). Topological domains in mammalian genomes identified by analysis of chromatin interactions. Nature 485, 376–380.

45. Moore, J.E., Purcaro, M.J., Pratt, H.E., Epstein, C.B., Shoresh, N., Adrian, J., Kawli, T., Davis, C.A., Dobin, A., Kaul, R., et al. (2020). Expanded encyclopaedias of DNA elements in the human and mouse genomes. Nature 583, 699–710.

46. Karp, N.A., Mason, J., Beaudet, A.L., Benjamini, Y., Bower, L., Braun, R.E., Brown, S.D.M., Chesler, E.J., Dickinson, M.E., Flenniken, A.M., et al. (2017). Prevalence of sexual dimorphism in mammalian phenotypic traits. Nature communications 8, 15475.

